# Motor-Like Properties of Non-Motor Enzymes

**DOI:** 10.1101/121848

**Authors:** David R. Slochower, Michael K. Gilson

**Affiliations:** Skaggs School of Pharmacy and Pharmaceutical Sciences, University of California San Diego, La Jolla, California 92093-0736, United States.

**Author notes:** Corresponding author: Michael K. Gilson, Skaggs School of Pharmacy and Pharmaceutical Sciences, University of California San Diego, La Jolla, California 92093-0736, United States.

**Keywords:** Molecular motors, protein dynamics, protein evolution

## Abstract

Molecular motors are thought to generate force and directional motion via nonequilibrium switching between energy surfaces. Because all enzymes can undergo such switching, we hypothesized that the ability to generate rotary motion and torque is not unique to highly adapted biological motor proteins, but is instead a common feature of enzymes. We used molecular dynamics simulations to compute energy surfaces for hundreds of torsions in three enzymes, adenosine kinase, protein kinase A, and HIV-1 protease, and used these energy surfaces within a kinetic model that accounts for intersurface switching and intrasurface probability flows. When substrate is out of equilibrium with product, we find computed torsion rotation rates up ~140 cycle s^-1^, with stall torques up to ~2 kcal mol^-1^ cycle^-1^, and power outputs up to ~50 kcal mol^-1^ s^-1^. We argue that these enzymes are instances of a general phenomenon of directional probability flows on asymmetric energy surfaces for systems out of equilibrium. Thus, we conjecture that cyclic probability fluxes, corresponding to rotations of torsions and higher-order collective variables, exist in any chiral molecule driven between states in a non-equilibrium manner; we call this the Asymmetry-Directionality conjecture. This is expected to apply as well to synthetic chiral molecules switched in a nonequilibrium manner between energy surfaces by light, redox chemistry, or catalysis.

## Introduction

A biological molecular motor is an enzyme that transduces chemical energy to mechanical motion. This motion must have a specific direction to fulfill the motor’s functional role. For example, a corkscrew-shaped flagellum must rotate in the appropriate sense to propel the organism. The ability to generate such directional motion appears to be a complex molecular property, perhaps unique to highly adapted motor proteins.

A system at equilibrium is guaranteed not to undergo net directional motion, due to the principle of microscopic reversibility. When the system is thrown out of equilibrium, there is the possibility of directional motion, but this requires a mechanism to capture and transduce the available chemical energy. Molecular motors work by using the chemical energy to drive stochastic switching between conformational energy surfaces in a manner that leads to directional motion (1-8). Non-motor enzymes also switch between at least two conformational free energy surfaces as they bind and release substrate. It is thus of interest to ask whether enzymes not normally thought of as motor proteins have motor-like properties when operating away from chemical equilibrium.

The present paper provides theoretical reasoning in support of this view, and then uses modeling to test the hypothesis that all periodic coordinates of any enzyme catalyzing an out-of-equilibrium reaction must undergo some degree of directional rotation. In order to uncover these rotations and begin to assess their magnitudes, we used molecular simulations and kinetic analysis to analyze the rotational dynamics of bond torsions induced by nonequilibrium, stochastic, state-switching in three, diverse, non-motor enzymes: adenosine kinase (ADK), protein kinase A (PKA), and HIV-1 protease (HIVP).

### Theoretical overview: asymmetric systems out of equilibrium

Consider a periodic molecular coordinate, θ, such as a bond torsion or a higher-order collective variable, in an enzyme undergoing a catalytic cycle of substrate-binding, catalysis, and product release. When binding of substrate switches the effective energy surface from that of the apo state, μ_0_(θ), to that of the bound state, μ_1_(θ), probability flows along θ towards a new equilibrium distribution corresponding to μ_1_(θ) (Figure 1). Such a flow of probability can be modeled with the Fokker-Planck equation, for example (4). Because enzymes are chiral, neither energy landscape, μ_0_(θ) and μ_1_(θ), can possess mirror symmetry; i.e., neither can be an exactly even function around any value of θ. Thus, there will be more probability flow in one direction than the other, due to the asymmetry of the starting probability distribution and of the patterns of barriers and troughs along the bound free energy surface. After the substrate is catalytically removed, this process repeats itself, with a probability distribution that has approached equilibrium on the substrate-bound surface diffusing asymmetrically under the potential of the apo free energy surface, μ_0_(θ). Unless there is an infinitely high energy barrier at the same location on both surfaces, there is no basis to expect that the net flows will cancel identically for a chiral molecule, and the resulting asymmetric net flow of probability during this two-step process constitutes directional motion. Note that any such motion will be superimposed on a background of rapid, non-directional stochastic fluctuations in θ.

**Fig. 1.**
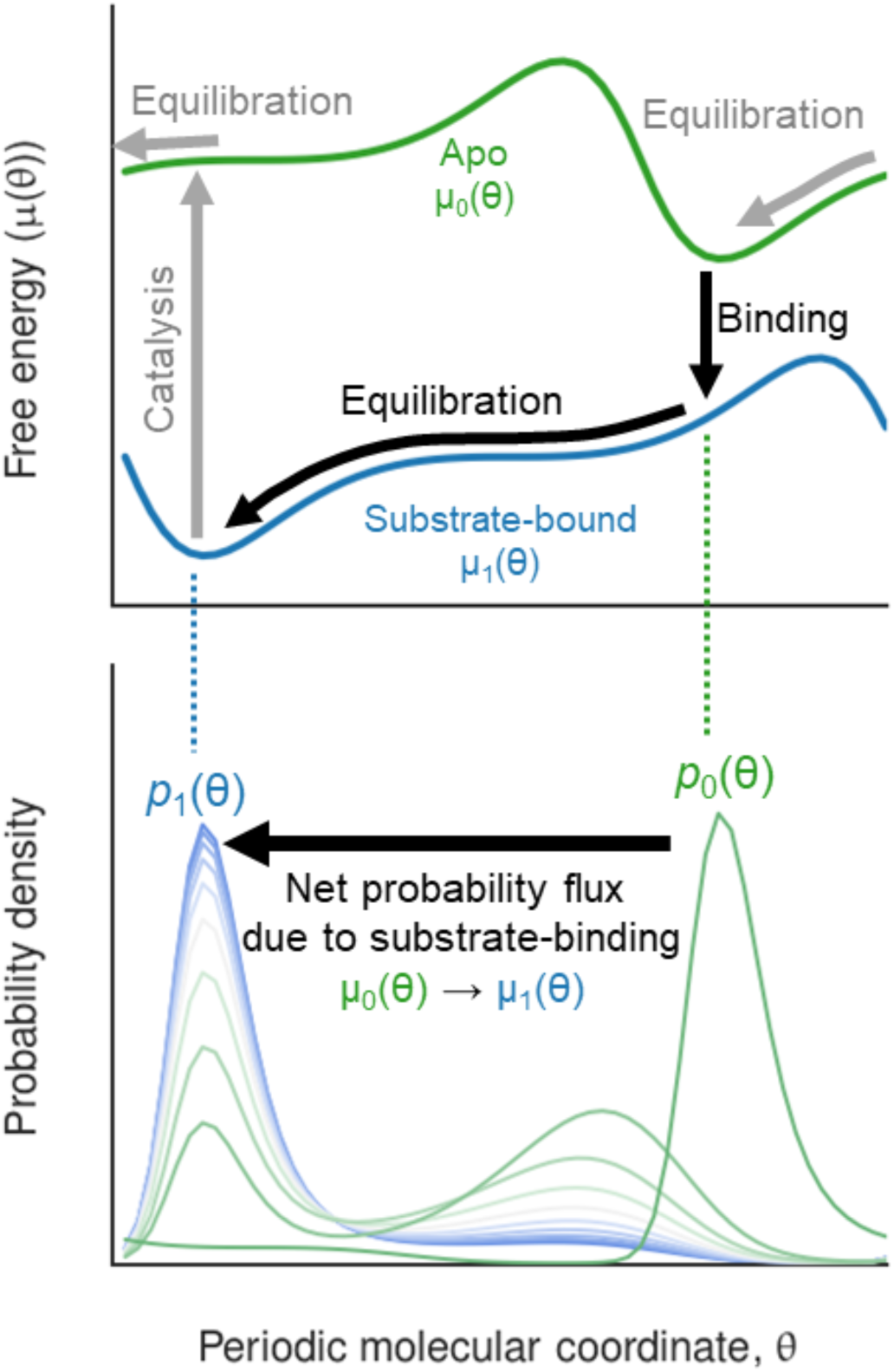
Illustrative apo and substrate-bound free energy surfaces of an enzyme (solid green and blue lines, respectively) and the flow of probability along coordinate θ, after the effective potential changes due to substrate binding (green to blue probability distribution curves).

## Materials and Methods

The one-dimensional free energy surfaces of protein main-and side-chain torsions, discretized into bins, were obtained from equilibrium molecular dynamics (MD) simulations of the enzymes in their apo and substrate-bound states (Supplementary Methods). These data, coupled with literature values for the enzyme kinetic parameters (Table S1), enabled us to define first order rate constants for transitions along and between the free energy surfaces (Fig. S1). The resulting set of rate equations was solved for the non-equilibrium steady state probability distribution, for a given concentration of substrate, and this, in turn, was used to compute the probability flux on each surface. The net flux, *J*, in units of torsional rotational cycles per second, is an indication of directional rotation; e.g., a positive value implies clockwise rotation of the torsion.

The three non-motor enzymes studied have distinctive characteristics: adenylate kinase (ADK), with 214 residues and a relatively high *k*_cat_ ~300 s^-1^ (9, 10), undergoes extensive conformational change on binding substrate, with two domains reorienting to form a compact conformation (11, 12); protein kinase A (PKA), with 350 residues and *k*_cat_ ~140 s^-1^ (13), acts as a “dynamic switch”, with long-range allosteric interactions and domain rearrangement upon ligand binding (14); and HIV-1 protease (HIVP), with 200 residues and lower *k*_cat_ ~10 s^-1^ (15-17), contains two flexible flaps that lose mobility in the substrate-bound state (18, 19) (Fig. S2).

## Results

For all three enzymes, we find that multiple side-chain and main-chain torsion angles undergo directional rotations, as indicated by nonzero probability flux, when excess substrate is present. Thus, at high substrate concentration, about 40 torsions in ADK and PKA rotate faster than 10 cycle s^-1^, and about 140 rotate faster than 1 cycle s^-1^ (Fig. 2a). Directional rotation is also observed for HIVP, though the rates are lower (Fig. 2b, red). The lower rates largely reflects the lower *k*_cat_ value of HIVP, relative to PKA and ADK. Thus, artificially assigning *k*_cat_ = 200 s^‒1^ to HIVP leads to substantial increases in the number of torsions with fluxes of at least 10 s^-1^ and at least 1 s^-1^ (Fig. 2b, orange and Fig. S4). The tendency toward lower fluxes in HIVP may also reflect the smaller scale of its conformational changes (Fig. S2): a small conformational change may lead to more similar energy surfaces in the two states, and hence less opportunity to generate rotational flux by the mechanisms discussed below. For all three enzymes, there are main chain torsions with substantial directional flux, but directional flux is more prevalent among side-chain torsions than main-chain torsions; see Fig. 3 and Fig. S3.

**Fig. 2.**
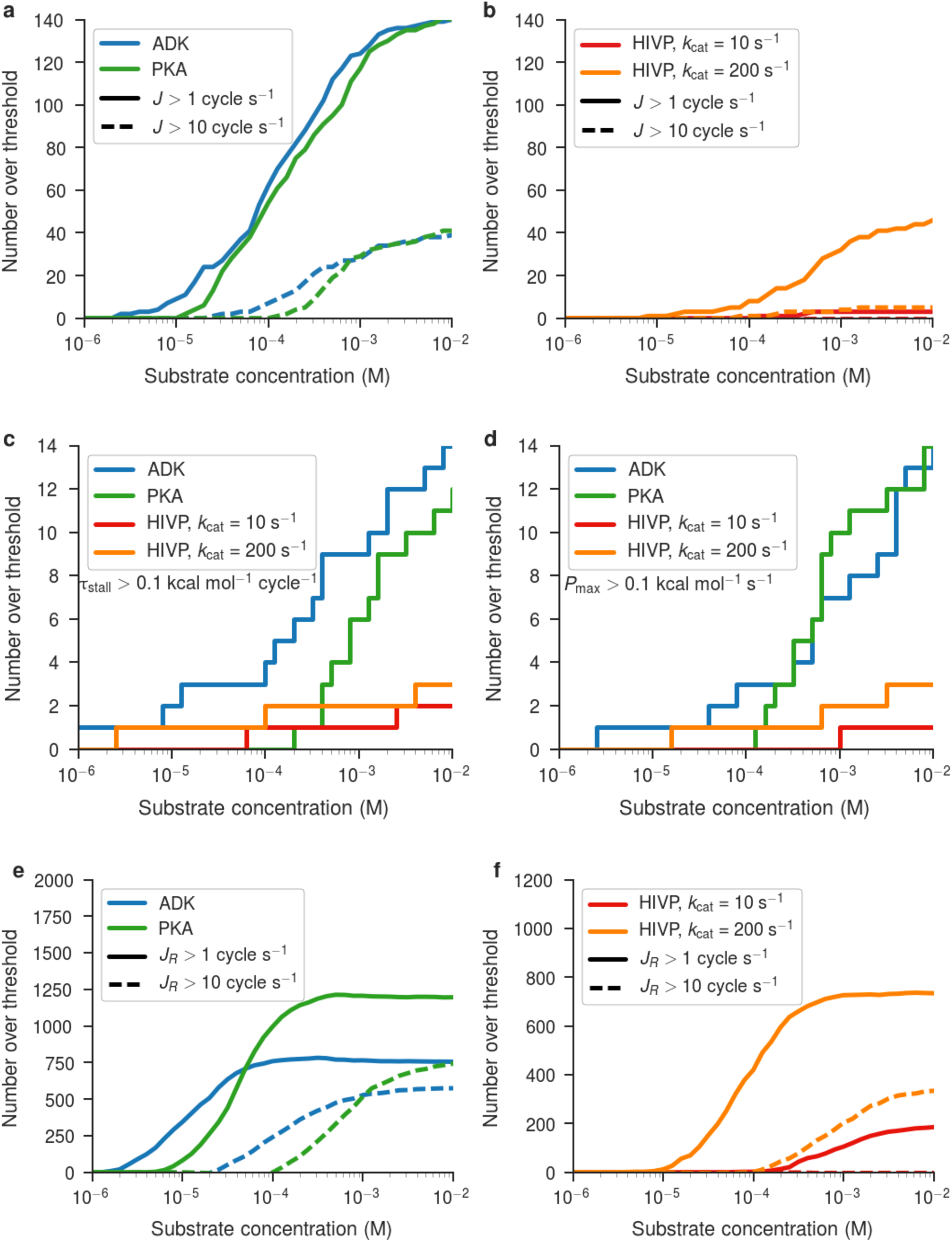
The number of torsions above various thresholds of directional flux magnitude, reciprocating flux magnitude, stall torque, and maximum power, as a function of substrate concentration. (a-b) The number of torsions with directional flux above 1 (solid) or 10 (dotted) cycle s^-1^ in ADK, PKA, and HIVP. (c-d) The number of angles with (c) maximum stall force above 0.1 kcal/(mol·cycle) and (d) power above 0.1 kcal/(mol·s^-1^). (e-f) The number of torsions with reciprocating flux above 1 (solid) or 10 (dotted) cycle s^-1^ and, at the same time, directional flux less than 1 cycle s^-1^.

**Fig. 3.**
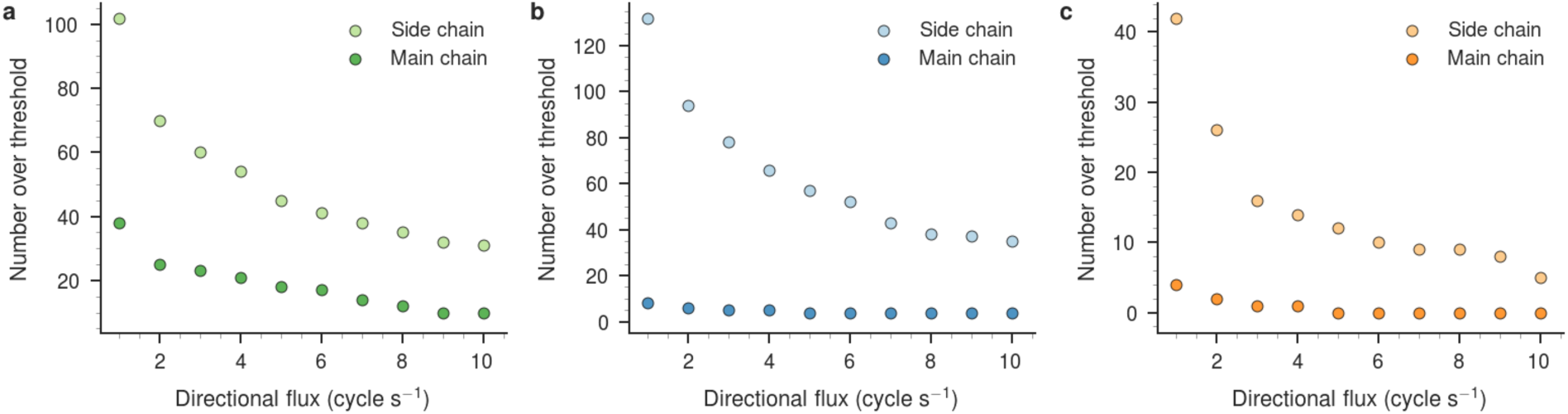
The number of main chain (ϕ and ψ) and side chain (χ) torsions above various thresholds of directional flux magnitude in each enzyme. Results are for (a) PKA, (b) ADK, and (c) HIVP (assuming *k*_cat_ = 200 s^-1^, see main text) at a substrate concentration of 10^-2^ M.

Although the maximum rotation rates are different for ADK, PKA and HIVP (180, 70 and 5.6 s^-1^, respectively), the maximum numbers of rotations per catalytic step are similar, at 0.5 – 0.6 cycles per catalytic turnover (Fig. 4). (Assuming *k*_cat_ = 200 s^-1^ for HIVP yields a maximum rate of 112 cycles s^-1^, which corresponds to 0.6 cycles per catalytic turnover.) This ratio is akin to a 2:1 gearing of catalysis to torsional rotation. Figs. S4 and S5 provide further details regarding the relationships between catalytic rate and torsional flux for additional torsions. The angles with highest directional flux are localized near the substrate binding pocket or mobile regions (Fig. S2, right two columns).

**Fig. 4.**
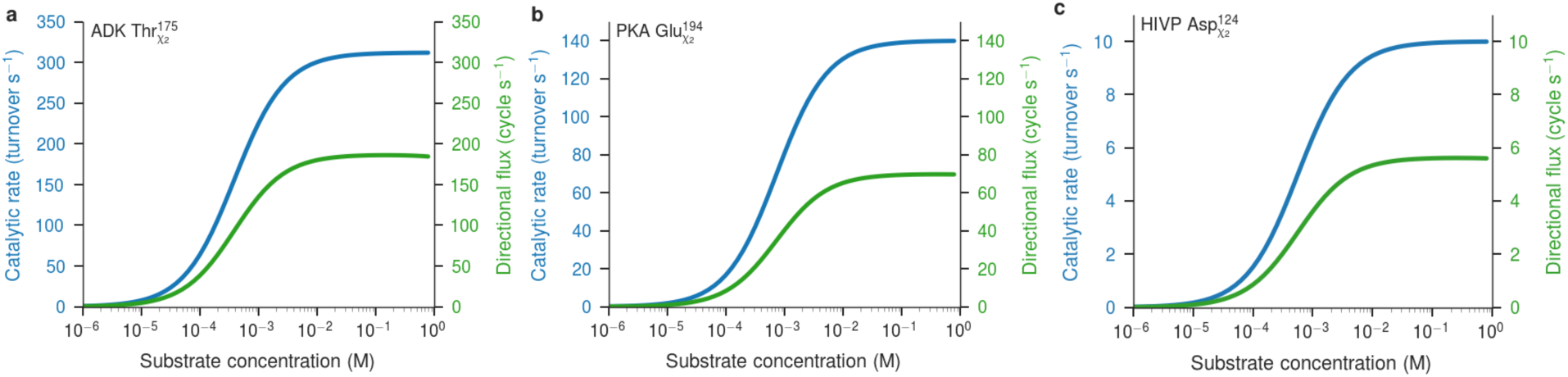
Dependence of catalytic rates and of the magnitude of directional flux on substrate concentration, for torsion angles in each enzyme. (a) The *χ*_2_ angle of Thr175 in ADK reaches a high level of directional flux. (b) The *χ*_2_ angle of Glu194 in PKA reaches a moderate level of directional flux. (c) Although the total amount of flux in the *χ*_2_ angle of Asp124 in HIVP is low, the ratio of directional flux to the enzyme velocity is similar to that in ADK and PKA.

We furthermore evaluated power output and performance under load by tilting the energy surfaces to generate a torque, *τ*, opposite to the directional flux, which modifies the intrasurface bin-to-bin rate constants (Supplementary Methods). The power output is the product of load torque and flux: *P* = *τJ*. Both the maximum power and the stall torque, *τ*_stall_, defined as the torque that brings the directional flux to zero, were found by scanning across values of applied torque. The results indicate that torsions in these enzymes can do work against small mechanical loads and thus generate power (Fig. 2c,d). In particular, at high substrate concentrations, torsions in ADK and PKA are predicted to generate stall torques up to 2.4 and 1.6 kcal mol^-1^ cycle^-1^, respectively, and maximum power outputs per torsion of 70 and 28 kcal mol^-1^ s^-1^.

The mechanism by which high rates of directional flux are generated is illustrated by the *χ*_2_ torsion of ADK Thr175 (Fig. 5a). This angle has a two-peaked probability distribution in both the bound and apo states, but the peak near 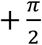 is favored in the apo state, while that near 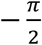 is favored in the bound state (Fig. 5b,c). In the presence of substrate, the bound-state energy minimum near 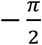 is highly occupied (Fig. 5b,c). Catalytic breakdown of substrate pumps the system to the secondary energy minimum of the apo state at 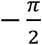 (Fig. 5c, arrow 1). Probability then flows primarily to the left on the apo surface, because this is the lowest-barrier path to the apo state’s global energy minimum near 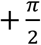 (arrow 2; this flux goes through the periodic boundary at *θ* = -*π* ≡ +*π*). Probability pooled in the global energy minimum of the apo state near 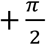, then flows primarily to the bound state, by binding substrate and landing in the secondary energy minimum of the bound state (arrow 3). It then flows back to the global minimum of the bound state via the lowest-barrier path, which is again leftward (arrow 4). The net effect is a leftward flux of up to −140 cycles s^-1^. Fig. 5d shows the steady state flux on each surface: leftward flux predominates overall, but occurs on the apo surface between 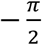 and 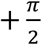, and on the bound surface elsewhere, with crossovers between surfaces at the energy minima.

**Fig. 5.**
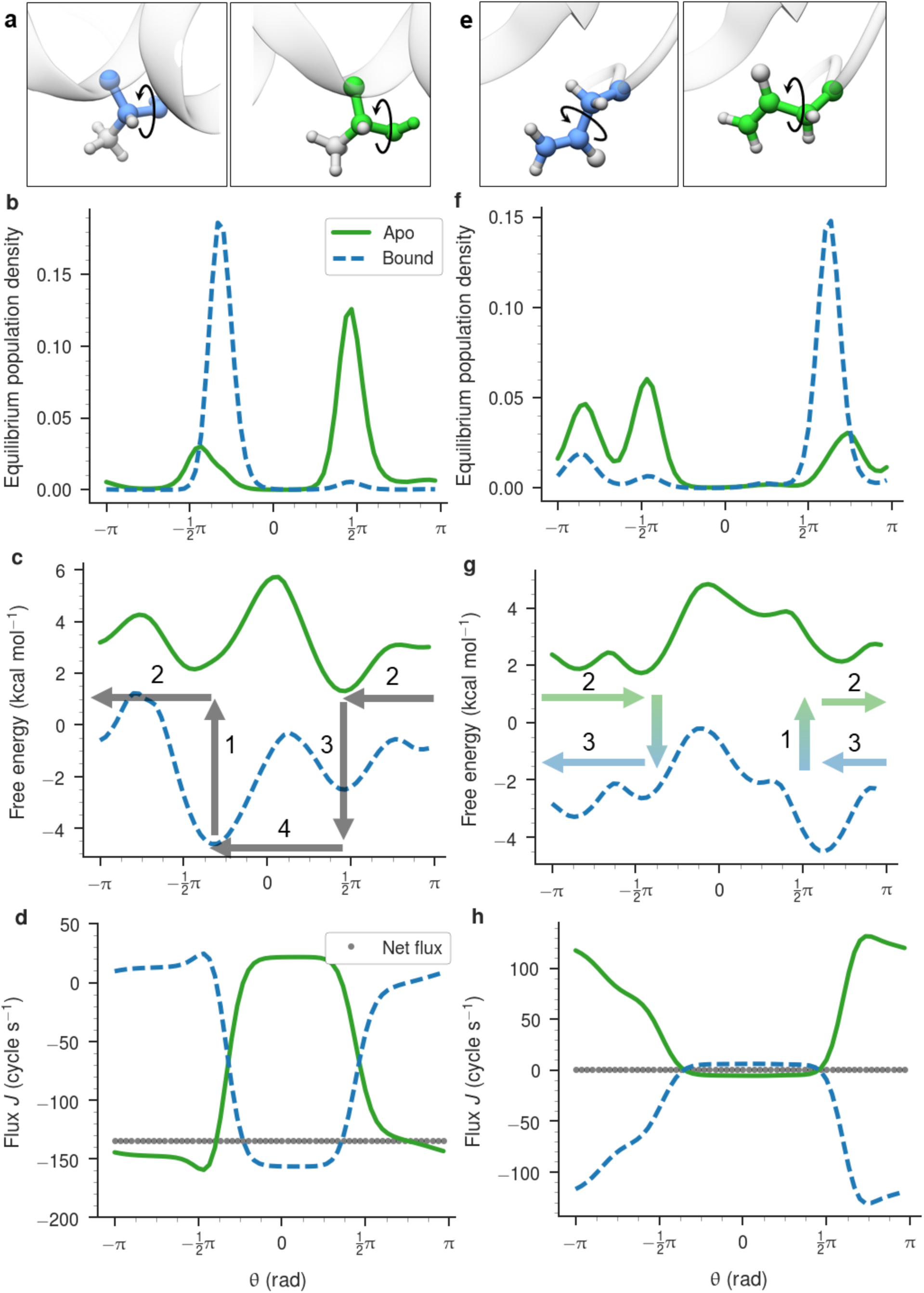
Protein torsion angles show directional and reciprocating motion. (a) ADK Thr175 in its crystallographic conformations for the apo (green) and bound (blue) forms (see Supplementary Methods for PDB accessions) with the *χ*_2_ angle denoted. The coloring is the same for panels a through d. (b) Equilibrium population densities of this angle from MD simulations (Supplementary Methods). (c) Free energy surfaces of this angle (Supplementary Methods) derived from the population densities in panel b. Arrows indicate the direction of probability flux along, and between, the two surfaces. (d) The probability flux drawn separately for each surface and as a sum (grey points), indicating large directional and reciprocating fluxes. (e-h) Same as a-d for ADK Asn138. In all cases the substrate concentration is 10^-3^ M.

This process parallels fluctuating potential mechanisms previously invoked to explain highly evolved molecular motors (3, 4, 7, 8, 20-23). Intuitively, high flux can be generated when each energy surface (apo and bound) has a main energy barrier and a main energy well, and the two surfaces are offset, so that probability flow in a given direction on each surface bypasses the barrier on the other surface, as schematized in Fig. S6 a-b. Also, as noted above, generation of net flux requires asymmetric energy surfaces; i.e., energy surfaces that are not even functions around any value of θ. Accordingly, symmetrizing the energy surfaces of a high-flux torsion abolishes directional flux, as shown in Fig. S7.

There are also many torsions whose net directional flux is small (< 1 cycle s^-1^), and whose dynamics are instead dominated by reciprocating motion, in which probability flows in one direction in the bound state and in the other direction in the apo state, within some angular range. These reciprocating probability fluxes correspond to driven, oar-like motion of the atoms controlled by the torsion. The enzymes ADK and PKA, respectively, are predicted to have ~1250 and ~750 torsions with reciprocating motions at rates of at least 1 cycle s^-1^ (Fig. 2e,f), and minimal directional flux. The maximal reciprocating fluxes are greater than the maximal directional fluxes, and, for ADK and PKA, approach the catalytic rates (Fig. S5). The mechanism of reciprocating motion is illustrated by *χ*_2_ of Asn 138 in ADK (Fig. 5e), which has a net flux that is nonzero but well below 1 cycle s^-1^, yet has reciprocating fluxes of up to 130 cycles s^-1^ (Fig. 5e-h). Reciprocating flux occurs when the main energy wells on the two surfaces are in different locations, but little probability flow on each surface bypasses the main barrier on the other surface, as schematized in Fig. S6 c-d. Thus, if the main barriers in the two states are coincident, movement will be dominated by reciprocating motion. It is worth noting that such reciprocating motion could generate net directional motion, if there were an additional mechanism that decoupled the load from the torsion for motion in either the bound or apo state. This would require correlated motion along at least one additional degree of freedom.

## Discussion

We have provided reasoning and evidence that directional rotation which can work against a load is not limited to highly adapted biological motor proteins. Instead, any enzyme operating out of equilibrium should generate directional conformational fluxes. The magnitudes of these motions will depend on a number of factors, including the levels and locations of the barriers and energy wells on the energy surfaces associated with different states of the system, and the chemical driving forces and kinetic constants. We also observe high reciprocating flux in torsions with energy surfaces that are not well tuned to generate net directional motion.

The present modeling approach uses atomistic simulations to obtain effective energy surfaces, and then embeds these in a kinetic model. This efficient technique is broadly applicable. For example, it could be used to study not only proteins but also synthetic molecular motors. It can also be extended to higher-dimensional distribution functions, and hence multidimensional energy surfaces, which will come into play when considering larger scale cooperative motions. Such motions could lead, for example, to state-dependent coupling of a load to a driven, reciprocating degree of freedom (see above). Note, however, that substantially more conformational sampling would be necessary to populate the multidimensional histograms arising in this application.

The basic mechanism by which directional motion is generated here is not limited to enzymes. We conjecture that the same statistical mechanical considerations will cause any system which undergoes stochastic, nonequilibrium switching between asymmetric energy surfaces to generate directional, cyclic motion. We term this idea the Asymmetry-Directionality (AD) conjecture. The AD conjecture applies to many molecular systems other than enzymes, such as ones where states are switched by absorption of light, changes in electrical potential, or changes in solution conditions. Thus, the occurrence of directional cycles of conformational change in molecular systems out of equilibrium should be regarded as the rule, rather than the exception.

The same reasoning applies not only to one-dimensional torsional motions, but also to protein motions associated with large-scale conformational change, because the associated multidimensional energy surfaces have asymmetric potentials, due to chirality. Such motions might help explain the observation that the translational diffusion coefficients of some non-motor enzymes rise with substrate concentration and hence with enzyme velocity (24, 25). The generality of the AD conjecture also means that enzymes from earliest evolutionary time would have had the ability to generate directional motion, and thus would have had a good starting point from which to embark on an evolutionary path to today’s highly adapted motor proteins.

## Acknowledgments

We thank Dr. N.-L. Huang for assistance preparing simulations on ADK and HIVP; Mr. M. Feng and Drs. A. Gilson, K. Lindenberg, C. Van den Broeck, and J.A. McCammon for theoretical discussions; and Dr. C. McClendon for providing the PKA simulations. This work was funded in part by grant GM061300 from the National Institutes of Health (NIH). Its contents are solely the responsibility of the authors and do not necessarily represent the official views of the NIH.

M.K.G. has an equity interest in and is a cofounder and scientific advisor of VeraChem LLC.

The datasets generated and analyzed during the current study are available in the GitHub repository https://github.com/GilsonLabUCSD/nonequilibrium.

M.K.G. conceived and designed the project. D.R.S. implemented the model and performed the simulations. M.K.G. and D.R.S. analyzed the data and wrote the manuscript.

## Supporting Material

### Fluctuating potential kinetic model

#### Definition of the model

The enzyme is considered to have two chief macrostates, apo (P_0_) or substrate-bound (P_1_), and is assumed to follow Michaelis-Menten kinetics:

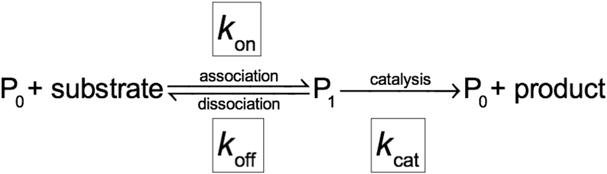

Thus, the substrate-bound protein can reach the apo state either through dissociation of the substrate or through catalytic conversion of the substrate into the product, followed by dissociation of the product. The substrate concentration is left as an adjustable parameter, while, the concentration of product is considered low enough that the reverse reaction makes no significant contribution to the kinetics.

The equilibrium probability distribution functions of the torsion angles of interest, in protein states *P*_0_ and *P*_1_ were obtained as follows. Torsional histograms, each with *N* bins, were computed from the equilibrium MD simulations described below, and converted to probabilities by normalization. The resulting probability distributions were smoothed with a Gaussian kernel (standard deviation of 1 bin), and any bins with zero probability after smoothing were assigned the lowest probability observed in any bin for that angle (typically about 10^-6^). Following normalization, the resulting probabilities are *p*_0,*i*_ and *p*_1,*i*_ for bin *i* in the apo and substrate-bound states, respectively, where 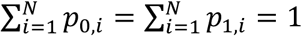. The present results are based on *N* = 60, which captures the shapes of the energy surfaces without imposing an excessive computational burden. Using fewer bins would effectively smooth the surfaces, reducing the differences between apo and bound and thus decreasing the computed fluxes and torques.

The kinetic model allows transitions between neighboring bins in the same state, with forward and reverse rate constants *k*_*x,i*→*i*+1_ and *k*_*x,i*+1→*i*_ respectively, where *x* = 0 for the apo state and *x* = 1 for the bound state; and transitions between states in the same bin, with rate constants *k*_0→1,*i*_ and *k*_1→0,*i*_ for binding and dissociation, respectively. These intersurface rate constants are set to

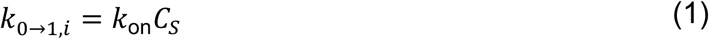

where *k*_on_ is the macroscopic on-rate for binding of substrate to the enzyme and *C*_*s*_ is the substrate concentration; and

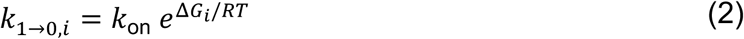

where Δ*G*_*i*_, the binding free energy associated with bin *i*, is given by

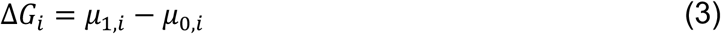

where *μ*_*x,i*_ is the free energy associated with bin *i* on surface *x*:

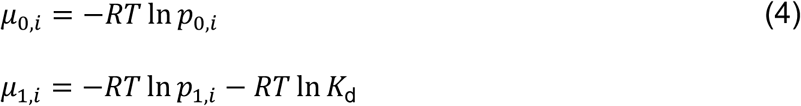

We now show that this set of rate constants leads to the correct dissociation constant, K_d_, for the full system. In particular, because *K*_d_ = *C*_1_ /*C*_0_*C*_*s*_, where *C*_1_ and *C*_0_ are the concentrations of bound and free protein, respectively, we want *p*_1_/ *p*_0_ = *C*_1_/*C*_0_ = *C*_*s*_ /*K*_d_, where *p*_0_ and *p*_1_ are the probabilities of finding the enzyme in the apo and bound states, respectively, so that *p*_0_ + *p*_1_ = 1. Using the fact that the equilibrium constant is the ratio of rate constants, and that the contribution of each bin’s rate constant to the corresponding macroscopic rate constant is weighted by the probability of the bin, we have

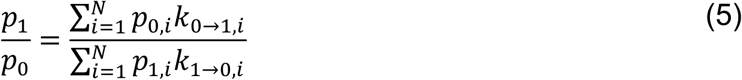

Using the prior equations, we obtain

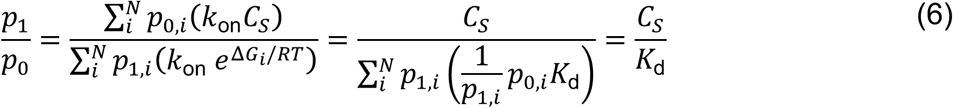

as required. This confirms that the intersurface rate constants in Eqs (1) – (2) correctly replicate the equilibrium between apo and bound enzyme in the absence of catalysis.

Transitions between neighboring bins on each energy surface are also modeled by first order rate processes, with the rate constants chosen to replicate free rotational diffusion in the limit of a flat energy surface, and to generate the MD-derived probability distribution on each surface in the absence of catalysis (see below):

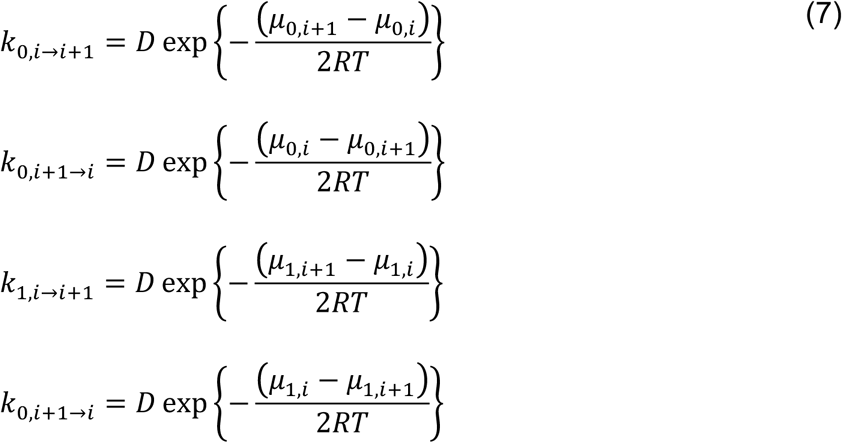

The forward and reverse rate constants are equal to *D* if the energy difference between the two bins is zero, and any energy difference increases and decreases the forward and reverse rate constants, respectively, by the same factor, as expected on physical grounds. The value of *D* is discussed in Assignment of Numerical Parameters, below.

Solving the first order kinetic system described above would yield an equilibrium Boltzmann probability distribution across the bins and states,

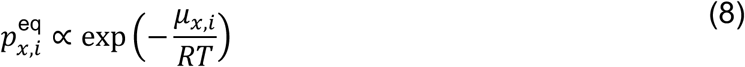

with the following normalization:

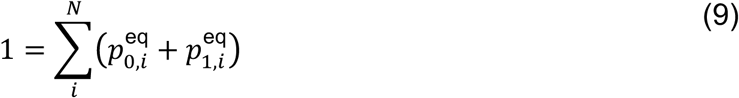

So far, we have included only one mechanism for transitioning from the bound state to the apo state: dissociative release of substrate. However, there is a second mechanism, which is the one that can put the system out of equilibrium, namely catalytic degradation of the substrate followed by product release. Referring to Eq (2), we account for this by adding another set of first-order rate processes going from the bound state bins to the apo state bins, with rate constants of *k*_cat_:

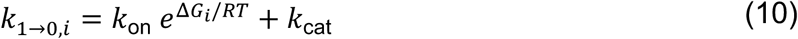

As noted above, for simplicity, we assume that the concentration of product is negligibly small. We also assume a uniform catalytic rate constant across bins, so that the ratio of binding to dissociation is *k*_0→1,*i*_/*k*_1→0,*i*_ = *k*_on_*C*_*s*_/(*k*_on_*e*^Δ*G*_*i*_/*RT*^ + *k*_cat_). Thus, high substrate concentrations drives substrate binding, while a favorable energy difference and/or a large catalytic rate constant drive the transition back to the apo state. When *C*_*s*_ and *k*_cat_ are nonzero, the system is no longer at equilibrium, but instead goes to a non-equilibrium, steady-state probability distribution, 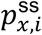. This distribution is obtained by the numerical method described in the following subsection.

#### Steady-state solution of the kinetic model

The system of rate equations is discretized in time by multiplying each rate constant by a small time-interval Δ*t*, yielding the fractional change in probability within each bin over this time. From these values, we construct a 2*N*×2*N* Markov transition matrix with off-diagonal elements that contain these fractional probability changes. The diagonal entries are then determined from the requirement that the sum of each row in the matrix equals 1. We used numerical methods (26, 27) to diagonalize the matrix, and the eigenvector whose eigenvalue value is unity corresponds to the steady state populations, 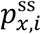 of all 2*N* bins in the system. The steady state probability flux between bins *i* and *i* + 1 on surface *x* is then

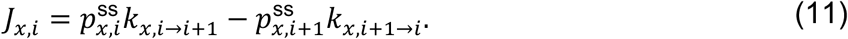

The net probability flux is the sum across both surfaces,

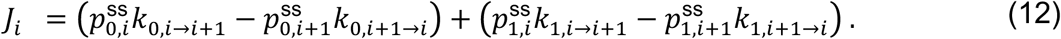

At steady state, *J*_*i*_ is uniform across bins, and we write *J* ≡ *J*_*i*_ in the main text as “directional flux”. We also report “reciprocating flux” for each torsion as the peak magnitude of directional flux across either surface, *J*_*R*_ = max(|*J*_0,*i*_|, |*J*_1,*i*_|).

#### Applying a load (torque) to a torsion angle

One test of these motors is their ability to do work against a mechanical load. We apply load by tilting the free energy surfaces with a constant slope, corresponding to a torque, τ, that opposes the direction of flux. This is done by supplementing the chemical potential differences in equation (4) with an energy difference Δ*E*, to generate rate expressions of the form

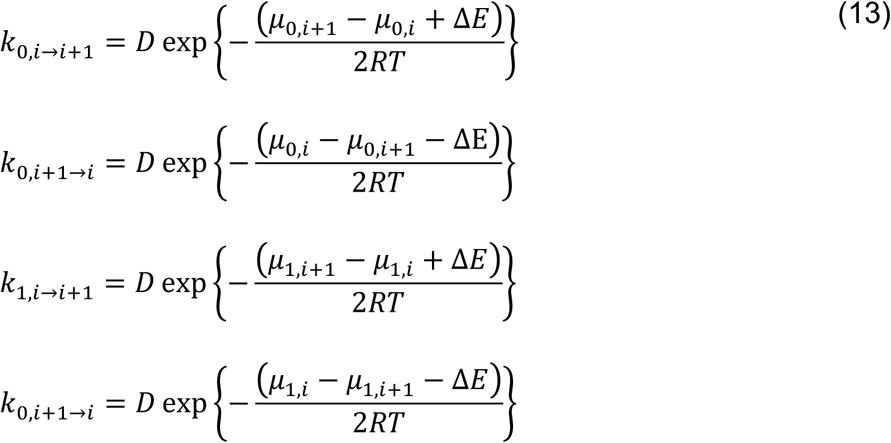

on both energy surfaces. That is, the energy difference between adjacent bins used to compute the transition rates (equation (9)) are modified by a factor ±Δ*E*, depending on direction, where Δ*E* > 0 gives a negative torque that is used to oppose positive probability flux, and vice versa. So that the load is correctly imposed across the periodic boundary, we replace *μ*_*x*,1_ – *μ*_*x,N*_ with *μ*_*x*,1_ – *μ*_*x,N*_ + Δ*E* and *μ*_*x,N*_ – *μ*_*x*,1_ with *μ*_*x,N*_ – *μ*_*x*,1_ – Δ*E*. The stall torque is the load that brings the next flux to zero; increasing the torque beyond this limit will cause the motor to slip backwards. We identify the stall torque by solving the kinetic system iteratively for a systematic scan of applied torques. The power produced for a given torque is given by *P* = *τJ*_*τ*_, where *J*_*τ*_ is the flux computed for a given applied torque τ. Empirically, the maximum power is found to occur at half the stall torque.

#### Assignment of numerical parameters

The present model requires numerical values of *D*, *k*_on_, *K*_d_ = 1/*K*_a_, and *k*_cat_. An initial estimate of *D* was made by simulating a molecule of butane, with the force field torsion and nonbonded terms set to zero to create a flat energy surface, and determining the rotational diffusion constant for this barrierless torsion. The mean value over one hundred 1 ns Langevin dynamics simulations is 3×10^15^ degree^2^ s^-1^. However, this large value led to numerical instability during diagonalization of the transition matrix, so we tested how the computed fluxes vary with D, and found that they are insensitive to the value of *D* once it is above ~ 10^9^ degree^2^ s^-1^. We used *D* = 3×10^12^ degree^2^ s^-1^ for the present calculations, as this is well into the regime where the results are independent of *D*, but not so large as to cause numerical problems (Fig. S7). (More sophisticated solvers might allow one to solve the system with higher values of D (28).) The enzyme kinetic parameters and their basis in the experimental literature are provided in Table S1.

## Materials and Methods

### Torsion potentials of mean force from molecular dynamics simulations

#### Adenylate Kinase

PDB accession 4AKE (29) was used as a starting structure for apo ADK; crystallographically ordered water molecules were retained. Hydrogens were added with pdb2pqr (30) at pH 7.0, bringing the net charge of the protein to −4. The substrate-bound protein was similarly modeled using PDB accession 3HPQ (31), which includes the ligand AP5A, a transition state analog, bound to the active site. AP5A carries a charge of −5 on the five phosphate groups, and the protein again carries a charge of −4 in the bound state. Partial atomic charges for AP5A were determined using the AM1-BCC method (32) in the antechamber program, and the remaining force field parameters were assigned from GAFF (33, 34). The apo and substrate-bound simulation systems were neutralized with 4 and 9 sodium ions, respectively. Both protein structures were solvated in a truncated octahedron with 12 Å padding.

Each system was energy-minimized for 20,000 steps, thermalized to 300 K over 1 ns, and equilibrated for 100 ns. Production simulations were then carried out for 1.0 μs using PME electrostatics with a 9 Å cutoff, and hydrogen mass repartitioning (35) with a 4 fs time step, using `pmemd.cuda.MPI` module of Amber 16 (36). Histograms (60 bins) of the torsion types listed in Table S2 were computed over the entire production simulation, using the cpptraj module (37) of Amber 16.

#### HIV Protease

PDB accession 1HHP (38) was used to model the apo structure of HIV-1 protease; crystallographically ordered waters were retained. To be consistent with Uniprot P03367, we made the following computational mutations: K14R, S37N, R41K, L63P, and I64V. Hydrogens were added with pdb2pqr at pH 7.0, bringing the net charge of the protein to +4. The substrate-bound protein was modeled using PDB accession 1KJF (39) with 10 residues of the Gag protein co-crystallized (RPGNFLQSRP; residues 443-452; fragment p1-p6 or SP2-p6). The following mutations were made to make the apo and bound structures consistent: K7Q and N25D. The charge of the bound protein and peptide was +6. Both models were solvated in a truncated octahedron with 12 Å padding and 4 or 6 chloride ions, respectively, added to neutralize the charge. The simulation procedures were identical to those for ADK.

#### Protein Kinase A

Simulations were carried out as described in reference (14).

### Numerical convergence

Convergence of the results was tested by using increasing amounts of simulation time to derive apo and bound state population histograms and then predicting the directional flux based on those histograms. For a sample angle shown in Fig. S8, after 500 ns, the calculated directional flux is very close to its value at 1000 ns, and the uncertainty in the flux value drops from about ±60 cycle s^-1^ to about ±1 cycle s^-1^ going from 500 ns to 1000 ns.

---

**Fig. S1.**
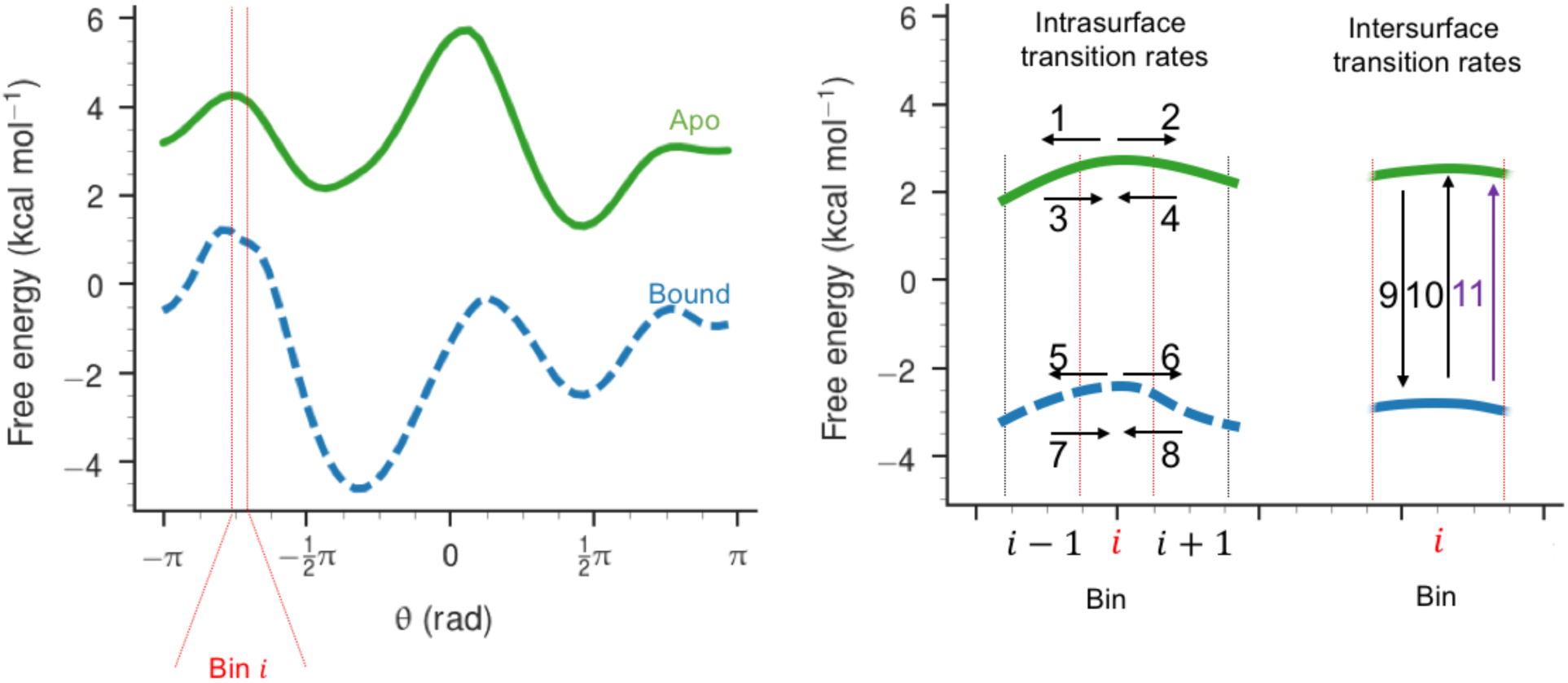
Transitions in the kinetic model. Small segments of the apo (green) and bound (blue) surfaces are diagrammed, with boundaries between discretization bins denoted with vertical lines. For each bin *i*, there are transitions in both directions between *i* and *i* – 1 (arrows 1 and 3) as well as between *i* and *i* + 1 (arrows 2 and 4) on the apo surface; there are similar transitions on the bound surface (arrows 5 and 7 and arrows 6 and 8, respectively). In addition, for each bin *i*, there is a transition from the apo surface to the bound surface (arrow 9, substrate binding) a dissociative transition from the bound surface to the apo surface (arrow 10), and a catalytic transition from the bound surface to the apo surface (arrow 11).

**Fig. S2.**
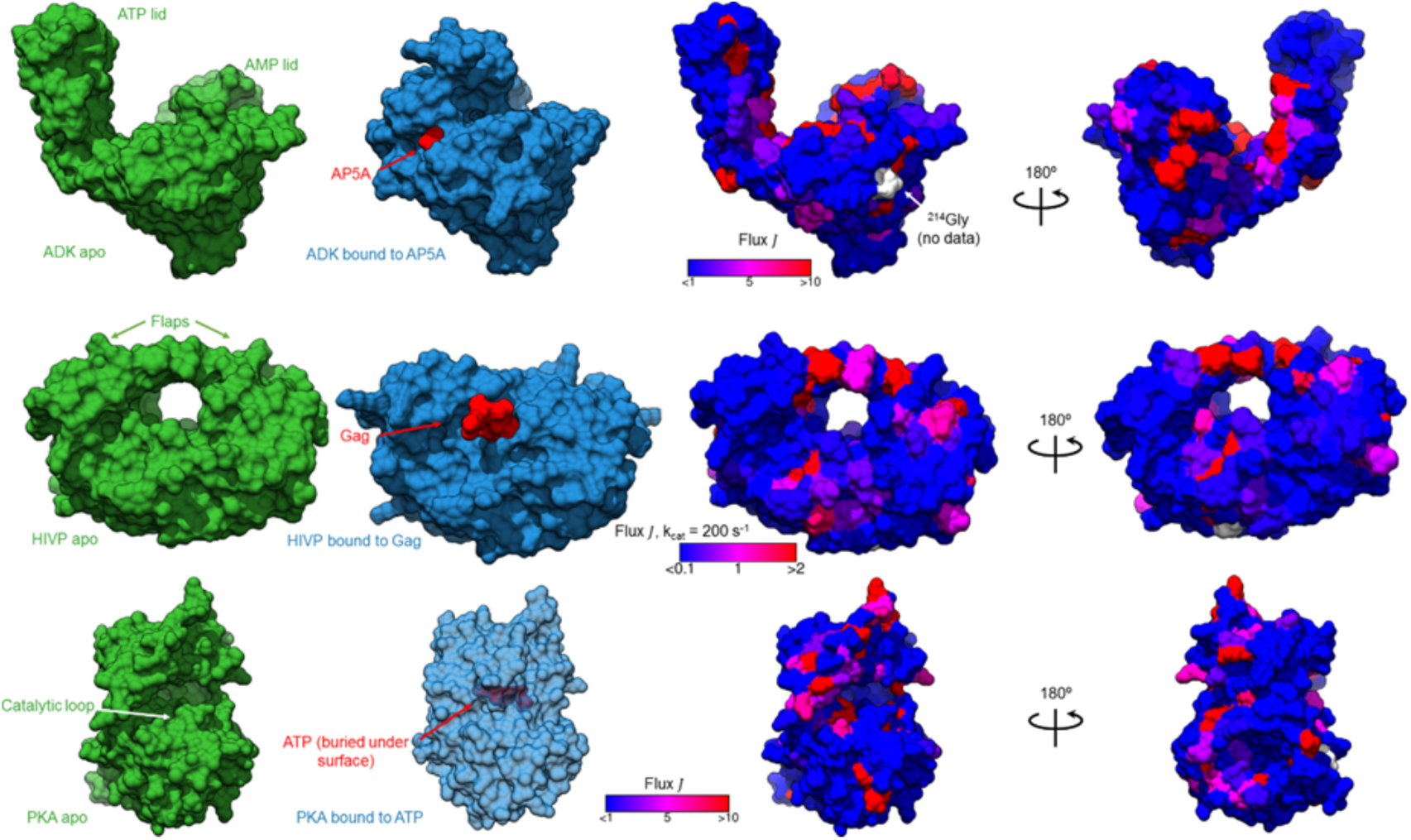
The apo and substrate-bound structures of the enzymes studied. Top row: the apo (green; PDB accession 4AKE) and substrate-bound (blue; PDB accession 3HPQ) conformations of ADK, rendered as a surface, with substrate AP5A (red) as spheres. In the right two panels, the absolute magnitude of directional flux *J* is mapped onto the apo structure, using a color gradient with thresholds from < 1 cycle s^-1^ (blue) to > 10 cycle s^-1^ (red). Middle row: the apo (green; PDB accession 1HHP) and substrate-bound (blue; PDB accession 1KJF) conformations of HIVP, rendered as a surface, with substrate Gag peptide (red) as spheres. In the right two panels, the absolute magnitude of directional flux *J* is mapped onto the apo structure, using the same color gradient, with thresholds < 0.1 cycle s^-1^ (blue) to > 2 cycle s^-1^ (red), to account for the lower level of directional flux in HIVP, even at k_cat_ = 200 s^-1^. Bottom row: the apo (green; PDB accession 1CMK) and substrate-bound (blue; PDB accession 3FJQ), with ATP (red) as spheres. In the right two panels, the absolute magnitude of directional flux *J* is mapped onto the apo structure, using the same thresholds as for ADK.

**Fig. S3.**
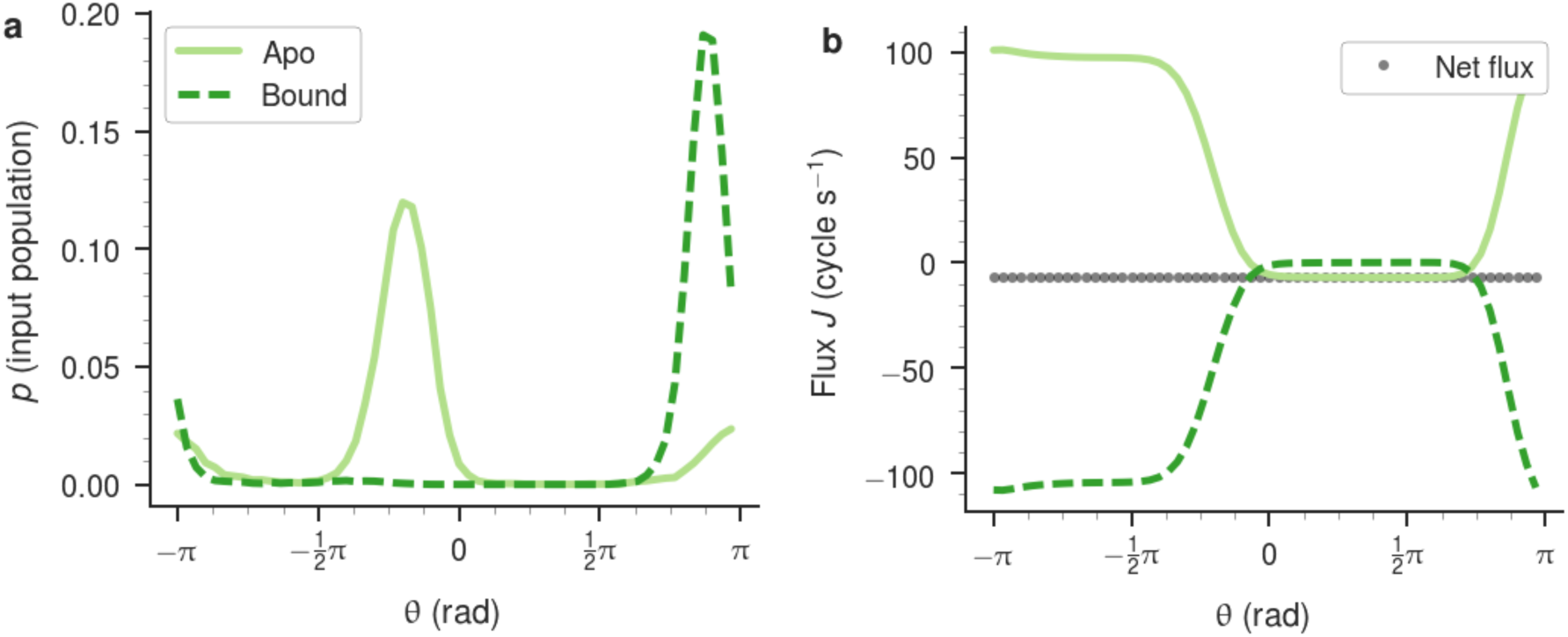
Ala110 ψ in PKA as an example of directional flux in a main chain torsion. (a) the population histograms and (b) −7 cycle s^-1^ net directional flux at a substrate concentration of 10^-2^ M.

**Fig. S4.**
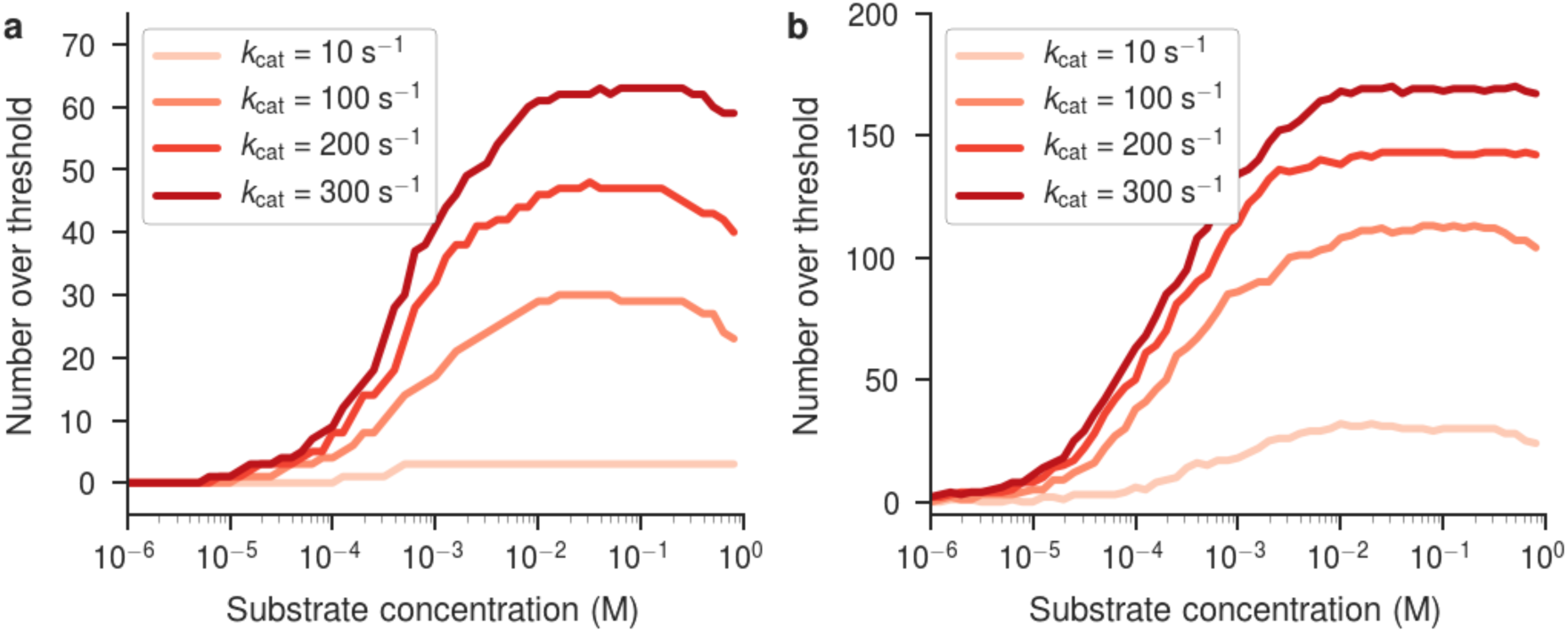
Number of HIVP torsion angles with net flux magnitude, |*J*|, above a given threshold, as a function of substrate concentration, for various assumed values of *k*_cat_. (a) The number of torsions with |*J*| > 1 cycle s^-1^ (b) The number of torsions with |*J*| > 0.1 cycle s^-1^.

**Fig. S5.**
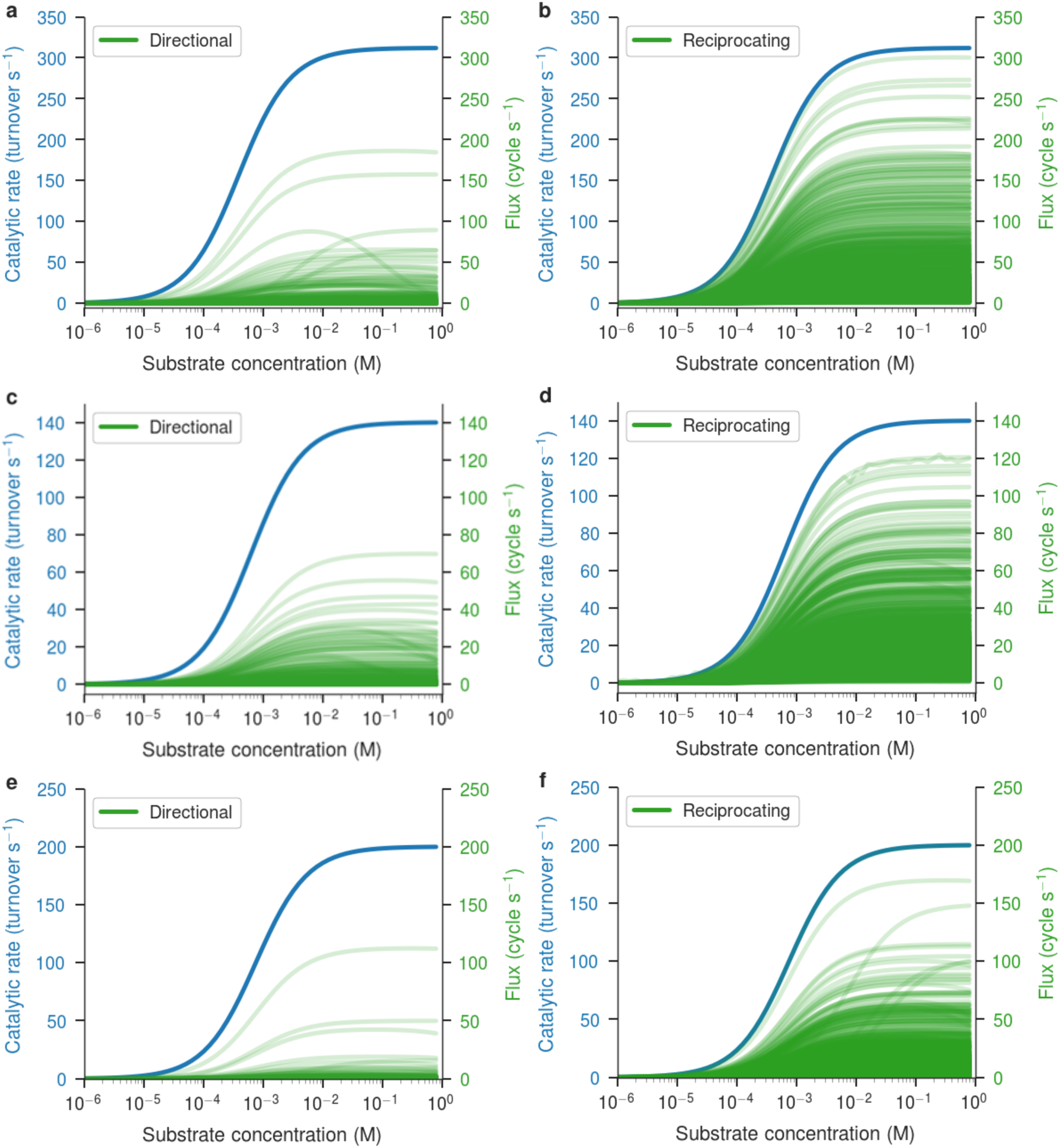
The dependence of the catalytic rate (blue) and flux (green) on substrate concentration. (a-b) All angles in ADK, calculated with *k*_cat_ = 312 s^-1^. (c-d) All angles in PKA, calculated with *k*_cat_ = 140 s^-1^. (e-f) All angles in HIVP, calculated with *k*_cat_ = 200 s^-1^. Reciprocating flux is shown only for angles with a maximum directional flux below 1 cycle s^-1^.

**Fig. S6.**
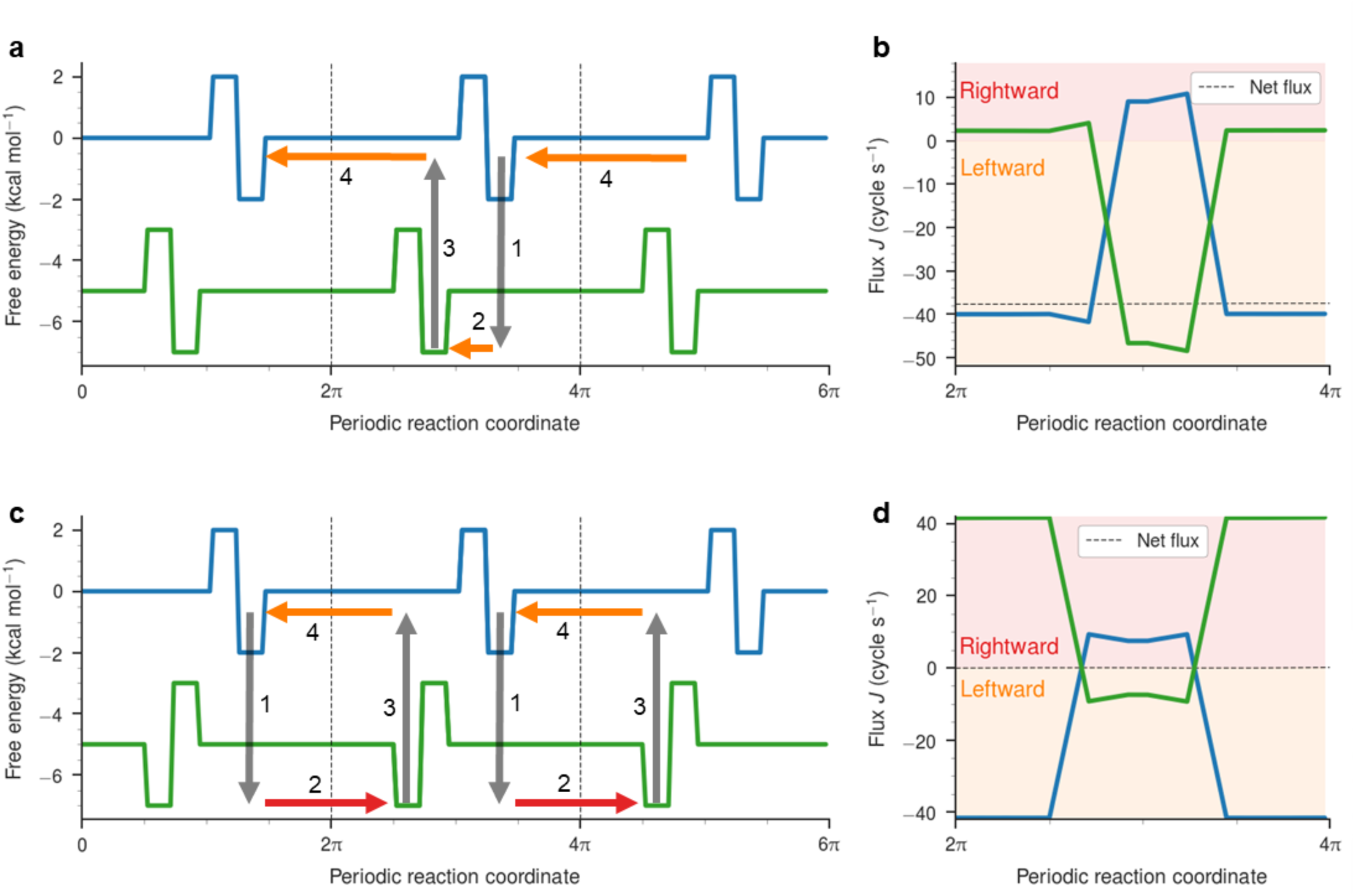
Sample periodic energy surfaces, each containing one barrier and one minimum per cycle, illustrating predominant directional or reciprocating net flux, depending on the relative positioning of the barriers and minima. (a-b) Energy surfaces that show net “left” directional flux *J*, when the probability is able to bypass the barrier on one surface by transitioning to the other surface. (c-d) Energy surfaces that show no directional net flux, but high reciprocating flux *J*_*R*_, arising from closed loops of probability flow.

**Fig. S7.**
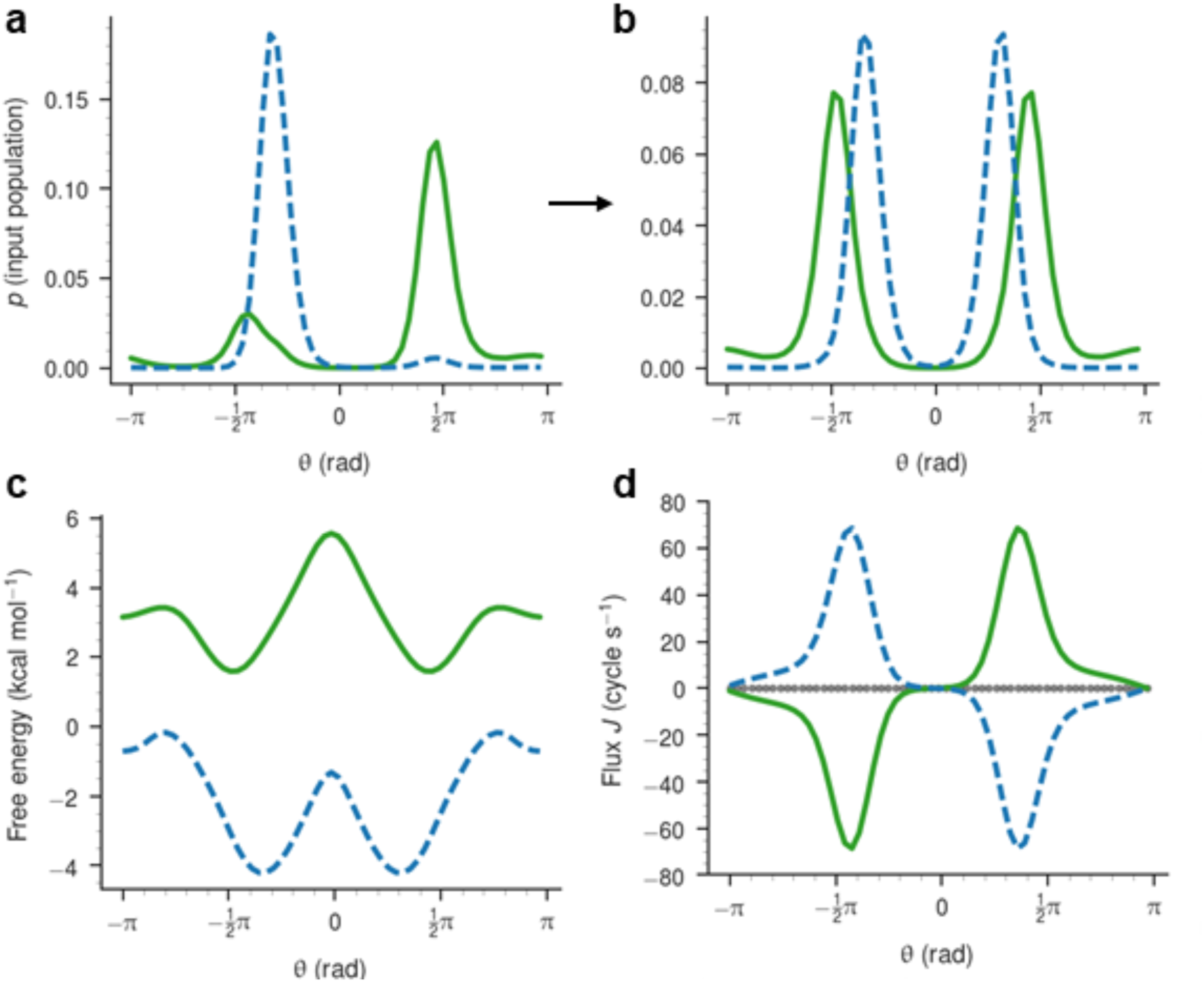
The effect of symmetrizing the energy landscape of the *χ*_2_ angle of ADK Thr175 on directional flux. (a) Unmodified probability distributions for the apo (green) and bound (blue) state of the torsion (also drawn in Fig. 3b). (b) The result of averaging each population histogram with its reflection about *θ* = 0 rad. (c) Free energy surfaces corresponding to the population densities in panel b (cf. Fig. 3c). The probability flux drawn separately for each surface and as a sum (grey points), showing exactly zero directional flux due the symmetry of the two energy surfaces (cf. Fig. 3d).

**Fig. S8.**
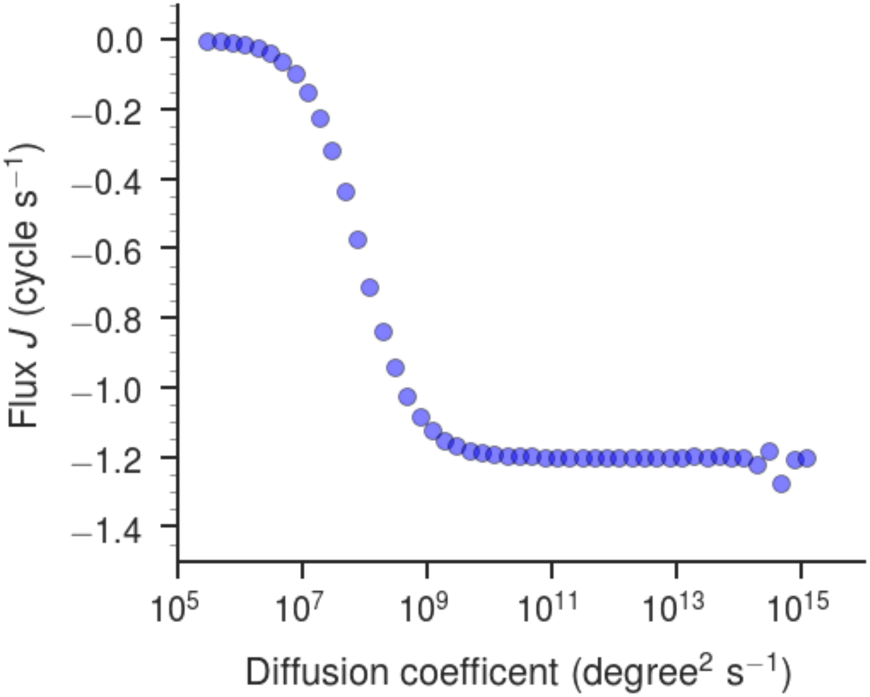
Flux as a function of the diffusion coefficient *D* for the *χ*_2_ angle of Thr175 in ADK.

**Fig. S9.**
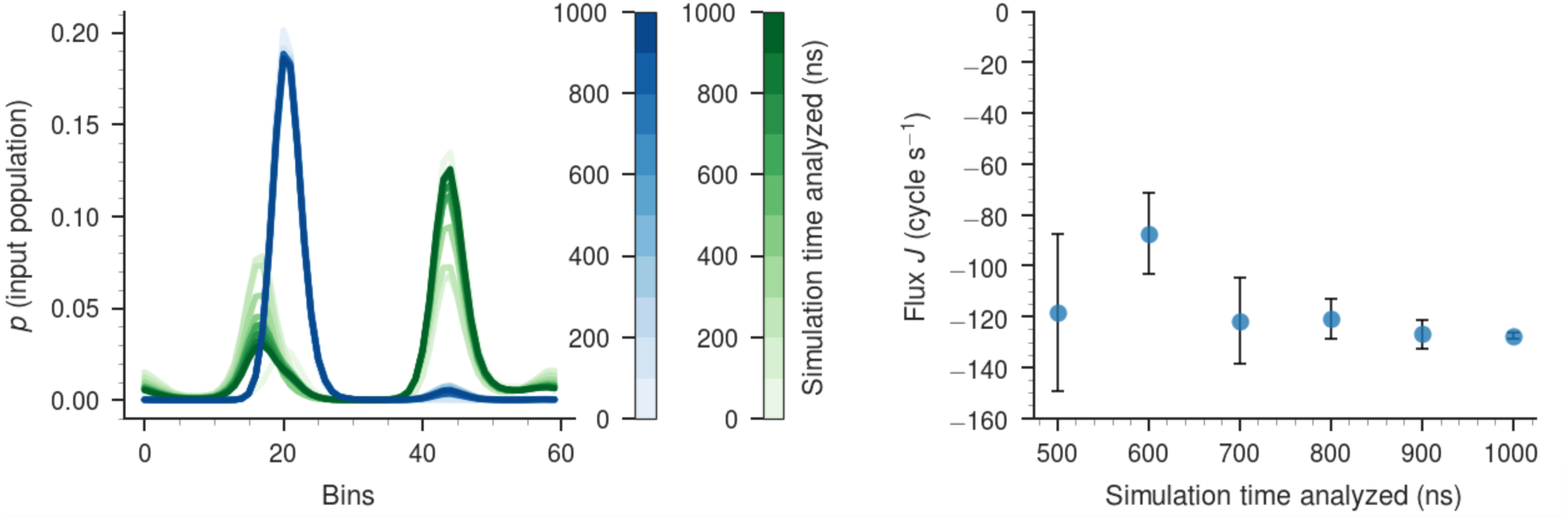
The convergence of the population histograms for the *χ*_2_ angle of Thr175 in ADK during equilibrium MD simulations for the apo (green) and bound (blue) states (left). The net flux calculated for this angle using the histograms on the left, after 500, 600, 700, 800, 900, and 1000 ns (right). The error bars represent the value of the net flux obtained after analyzing only the first or second half of each interval.

**Table S1.**
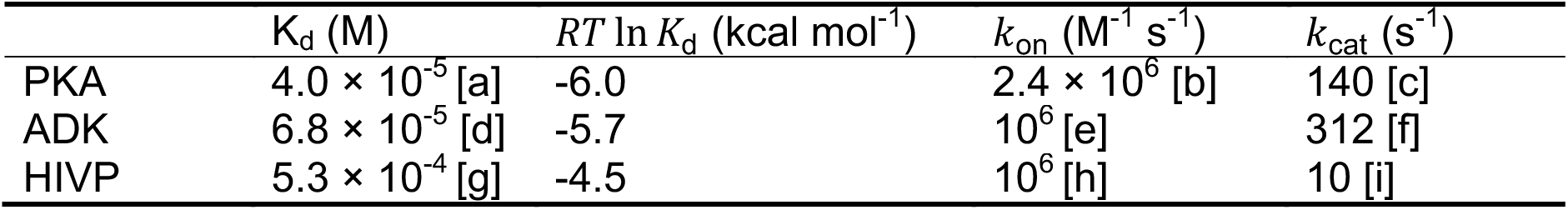
Values of enzymatic parameters used in the present calculations.

a. K_d_ for ATP in Scheme 4 in (40).
b. k_1_ in Scheme 4 in (40).1xA9k
c. The kinetics of PKA are complex and depend on, among other factors, the presence of divalent ion species and occupancy in the active site. The observed catalytic rate includes two or more conformational changes and lies between 50 s^-1^ and a fast, burst-phase of 500 s^-1^. We used the rate-limiting step (ADP · P_i_ release) from Figure 11 of (13) in this manuscript.
d. An average of K_d_ values for ATP found in (12) and (41). The calculated *μ*_offset_ is an average of the *μ*_offset_ for each K_d_. The K_d_ values for AMP are roughly four times as large. Our model implicitly assumes a single substrate and single on rate.
e. An order of magnitude estimate calculated using K_M_ from (9), k_cat_ values from (9) and (10), with the median expected ATP k_off_ from (12), and using the relationship 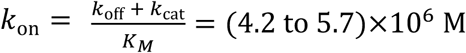, assuming Michaelis-Menten kinetics.
f. An average of k_cat_ from (9) and (10).
g. A literature value of K_d_ for the specific Gag fragment used in the simulations was not found in the literature. However, reference (42) reports K_M_ = 5.3 × 10^-4^ M for the p1/p6 substrate sequence PGNFLQS (the simulated bound peptide sequence was RPGNFLQSRP) and for sufficiently small catalytic rate, *K*_d_~*K*_M_. Several other references list values for k_cat_ / K_M_.
h. No value could be found in the literature, so the same order of magnitude as ADK is used.
i. Catalytic rates as low as 0.3 s^-1^ are reported in (42) for the p1/p6 substrate sequence, but other Gag sequences have catalytic rates on the order of 10 s^-1^, see for example (15-17).

**Table S2.** Torsion angle definitions, using atom types from Amber ff14SB.

| Residue | Angle | Atom 1 | Atom 2 | Atom 3 | Atom 4 |
| --- | --- | --- | --- | --- | --- |
| ALA | phi | C | N | CA | C |
| ALA | psi | N | CA | C | N |
| ALA | chi1 | HB1 | CB | CA | N |
| ARG | phi | C | N | CA | C |
| ARG | psi | N | CA | C | N |
| ARG | chi1 | CG | CB | CA | N |
| ARG | chi2 | CD | CG | CB | CA |
| ARG | chi3 | NE | CD | CG | CB |
| ARG | chi4 | CZ | NE | CD | CG |
| ARG | chi5 | NH1 | CZ | NE | CD |
| ASN | phi | C | N | CA | C |
| ASN | psi | N | CA | C | N |
| ASN | chi1 | CG | CB | CA | N |
| ASN | chi2 | ND2 | CG | CB | CA |
| ASP | phi | C | N | CA | C |
| ASP | psi | N | CA | C | N |
| ASP | chi1 | CG | CB | CA | N |
| ASP | chi2 | OD2 | CG | CB | CA |
| CYS | phi | C | N | CA | C |
| CYS | psi | N | CA | C | N |
| CYS | chi1 | SG | CB | CA | N |
| CYS | chi2 | HG | SG | CB | CA |
| GLN | phi | C | N | CA | C |
| GLN | psi | N | CA | C | N |
| GLN | chi1 | CG | CB | CA | N |
| GLN | chi2 | CD | CG | CB | CA |
| GLN | chi3 | NE2 | CD | CG | CB |
| GLN | chi4 | HE21 | NE2 | CD | CG |
| GLU | phi | C | N | CA | C |
| GLU | psi | N | CA | C | N |
| GLU | chi1 | CG | CB | CA | N |
| GLU | chi2 | CD | CG | CB | CA |
| GLU | chi3 | OE1 | CD | CG | CB |
| GLY | phi | C | N | CA | C |
| GLY | psi | N | CA | C | N |
| HIS | phi | C | N | CA | C |
| HIS | psi | N | CA | C | N |
| HIS | chi1 | CG | CB | CA | N |
| HIS | chi2 | CD2 | CG | CB | CA |
| ILE | phi | C | N | CA | C |
| ILE | psi | N | CA | C | N |
| ILE | chi1 | CG1 | CB | CA | N |
| ILE | chi2 | CD1 | CG1 | CB | CA |
| LEU | phi | C | N | CA | C |
| LEU | psi | N | CA | C | N |
| LEU | chi1 | CG | CB | CA | N |
| LEU | chi2 | CD1 | CG | CB | CA |
| LYS | phi | C | N | CA | C |
| LYS | psi | N | CA | C | N |
| LYS | chi1 | CG | CB | CA | N |
| LYS | chi2 | CD | CG | CB | CA |
| LYS | chi3 | CE | CD | CG | CB |
| LYS | chi4 | NZ | CE | CD | CG |
| MET | phi | C | N | CA | C |
| MET | psi | N | CA | C | N |
| MET | chi1 | CG | CB | CA | N |
| MET | chi2 | SD | CG | CB | CA |
| MET | chi3 | CE | SD | CG | CB |
| PHE | phi | C | N | CA | C |
| PHE | psi | N | CA | C | N |
| PHE | chi1 | CG | CB | CA | N |
| PHE | chi2 | CD1 | CG | CB | CA |
| PRO | phi | C | N | CA | C |
| PRO | psi | N | CA | C | N |
| PRO | chi1 | CG | CB | CA | N |
| PRO | chi2 | CD | CG | CB | CA |
| PRO | chi3 | N | CD | CG | CB |
| SER | phi | C | N | CA | C |
| SER | psi | N | CA | C | N |
| SER | chi1 | OG | CB | CA | N |
| THR | phi | C | N | CA | C |
| THR | psi | N | CA | C | N |
| THR | chi1 | CG2 | CB | CA | N |
| THR | chi2 | HG1 | OG1 | CB | CA |
| TRP | phi | C | N | CA | C |
| TRP | psi | N | CA | C | N |
| TRP | chi1 | CG | CB | CA | N |
| TRP | chi2 | CD1 | CG | CB | CA |
| TYR | phi | C | N | CA | C |
| TYR | psi | N | CA | C | N |
| TYR | chi1 | CG | CB | CA | N |
| TYR | chi2 | CD1 | CG | CB | CA |
| TYR | chi3 | HH | OH | CZ | CE1 |
| VAL | phi | C | N | CA | C |
| VAL | psi | N | CA | C | N |
| VAL | chi1 | CG1 | CB | CA | N |

